# Distinctive neural correlates of phonological processing and reading impairment in fetal alcohol-exposed adolescents with and without facial dysmorphology

**DOI:** 10.1101/2021.08.04.454704

**Authors:** Xi Yu, Jade Dunstan, Sandra W. Jacobson, Christopher D. Molteno, Nadine M. Lindinger, Theodore K. Turesky, Ernesta M. Meintjes, Joseph L. Jacobson, Nadine Gaab

**Author notes:** Corresponding authors: Xi Yu, PhD, State Key Laboratory of Cognitive Neuroscience and Learning, Beijing Normal University, Beijing 100875, China,; Nadine Gaab, PhD, Harvard Graduate School of Education, Cambridge, MA 02138, USA,; Joseph L. Jacobson, PhD, Wayne State University School of Medicine, Detroit, MI 48201, USA.

## Abstract

**Background:** Prenatal alcohol exposure (PAE) has been linked to atypical brain development and a wide range of cognitive and behavioral impairments, including poor reading performance in childhood and adolescence. However, little is known about how structural and/or functional teratogenesis in the brain mediate reading impairment in fetal alcohol spectrum disorders (FASD) and whether neural correlates of reading and phonological processing differ between FASD subtypes with different clinical presentations in facial morphology.

**Methods:** The current study utilized functional magnetic resonance imaging (fMRI) and diffusion tensor imaging (DTI) to characterize functional and structural mechanisms mediating reading deficits in 26 syndromal adolescents with PAE-related facial dysmorphology (i.e., fetal alcohol syndrome (FAS) or partial FAS (PFAS)) and 30 heavily exposed (HE) without this dysmorphology, in comparison with 19 typically developing controls. Importantly, the levels of PAE and reading ability were comparable between the FAS/PFAS and HE groups in the current study.

**Results:** Compared to the nonsyndromal HE and control groups, the syndromal adolescents showed greater activation in the right precentral gyrus during an fMRI phonological processing task and rightward lateralization in an important reading-related tract (inferior longitudinal fasciculus, ILF), suggesting an atypical reliance on the right hemisphere during reading. By contrast, in the HE group, better reading skills were associated with increased neural activation in the left angular gyrus (LAG) and higher fractional anisotropy in the white matter organization of the left ILF. However, the brain function-behavior relation was weaker in the HE than among the controls, suggesting less efficient function of the typical reading neural network that may contribute to the observed reading impairments.

**Conclusions:** Our findings provide the first evidence for the distinctive functional and structural mechanisms underlying atypical reading and phonological processing in PAE adolescents with and without FAS facial dysmorphology.

**Highlights:**

- Prenatal alcohol exposure is associated with altered neural reading networks
- FASD subtypes exhibit distinctive neural correlates of phonological processing
- Greater right-hemispheric reliance was observed in FASD with facial dysmorphology
- Non-syndromal FASD showed deficits in the typical left-hemispheric reading network

## 1. Introduction

Fetal alcohol spectrum disorders (FASD) refer to a wide range of physical, cognitive and behavioral impairments resulting from prenatal alcohol exposure (PAE; Glass et al., 2017; J. L. Jacobson et al., 2011; S. W. Jacobson et al., 2004; Mattson et al., 2019). Although these disorders frequently go undiagnosed due to social stigma, diagnostic complexity and referral bias (McLennan, 2015), FASD affect an estimated 1.1% to 5% of the U.S. and European population (May et al., 2018; Wozniak et al., 2019) and can range as high as 30.6% in some communities in South Africa (May et al., 2021). The most severe cases, which are diagnosed with fetal alcohol syndrome (FAS, Hoyme et al., 2005), show distinctive craniofacial dysmorphology, growth restriction, and small head circumference. A diagnosis of partial FAS (PFAS) is given to individuals who exhibit some but not all of the facial dysmorphology and growth restriction seen in FAS. Nevertheless, regardless of the presence of dysmorphology, cognitive and behavioral impairments are seen in all children with FASD, including heavily exposed (HE) individuals who lack the characteristic facial anomalies (e.g., Mattson et al., 2019).

Children with FASD often show poorer academic performance in both reading and arithmetic tasks, compared to their peers (Hoyme et al., 2016; Sampson et al., 1989; Streissguth et al., 1990). Several studies have documented mathematical deficits, such as number estimation (Kopera-Frye et al., 1996), beginning as early as infancy (Berger et al., 2019), and calculation (J. L. Jacobson et al., 2011) in both children and adults with FASD (Burden et al., 2005; Chiodo et al., 2004; Coles et al., 1991; Streissguth et al., 1990), even after control for IQ (Goldschmidt et al., 1996; Kerns et al., 1997). These deficits have been demonstrated using behavioral, ERP, and functional magnetic resonance imaging (fMRI) methodologies (e.g., Meintjes et al., 2010; Santhanam et al., 2009; Ben-Shachar et al., 2020). Although reading has been less extensively investigated, children and adolescents with PAE have shown poorer performance on standardized word reading and spelling tests, when compared to controls (Glass et al., 2015, 2017; Howell et al., 2006; Lindinger et al., 2018; Sowell, et al., 2008), and were more likely to fail to attain reading “benchmarks” in school in a large population sample (*N* = 4056, O’Leary et al., 2013).

While previous studies have examined cognitive mechanisms underlying reading deficits associated with PAE, application of neuroimaging techniques has the potential to unravel the neural underpinnings of these impairments. For example, impairment in phonological processing, a foundational literacy skill referring to the ability to recognize and manipulate the sound structure of words, has been observed in children with FASD (Adnams et al., 2007; Korkman et al., 2003; Streissguth et al., 1994). Moreover, these deficits have been shown to account for their poorer reading and spelling abilities compared to controls (Glass et al., 2015), suggesting that one of the potential cognitive etiologies of the PAE-associated reading impairments may lay in deficient phonological skills. At the neural level, typical reading development relies primarily on a left-lateralized network (Richlan et al., 2009, 2011, 2013) that comprises the temporoparietal region responsible for speech perception and phonological processing (e.g., Cattinelli et al., 2013; Pugh et al., 2001; Schlaggar & McCandliss, 2007), the temporooccipital cortex underlying the orthographic representation (e.g., Cohen et al., 2000; Vinckier et al., 2007), as well as the anterior frontal associated with syntactic and semantic processing and domain-general functions, such as, executive function and working memory (e.g., Bonhage et al., 2015; Fiez et al., 1996; Malins et al., 2016; Pugh et al., 2001; Rimrodt et al., 2009; Rodd et al., 2015). Individuals with FASD, however, exhibit atypical structural morphometries, including reductions in regional brain volumes and alterations in cortical gyrification (De Guio et al., 2014; Sowell et al., 2001, 2002), as well as disrupted developmental trajectories (Lebel et al., 2012) in the left posterior temporoparietal cortex, aligning with the atypical phonological skills frequently reported in individuals with FASD. Moreover, white matter tracts underlying the typical reading network (McCandliss & Noble, 2003; Ozernov-Palchik & Gaab, 2016; Pugh et al., 2000) have been shown to be altered in FASD (Donald et al., 2015; Fan et al., 2016; Lebel et al., 2008; Sowell, et al., 2008). PAE-associated changes in the structural asymmetries of the temporal lobes have also been reported (Sowell, et al., 2002). These findings suggest that structural brain characteristics important for reading and its sub-components may be particularly susceptible to alcohol teratogenesis. However, empirical evidence is lacking as to whether and how the functional alterations observed in children with FASD are associated with their deficits in reading and its sub- components, such as phonological skills.

Another important question is whether distinctive neural characteristics of atypical cognitive function, reading and foundational literacy skills in this case, might emerge in FASD subtypes that differ in presentation of facial morphology (Moore et al., 2014). Significant correlations have been observed between PAE-related facial morphologies and brain volumes (Lebel et al., 2012; Roussotte et al., 2011), suggesting a link between brain structure and facial dysmorphology. Moreover, differences in PAE-associated brain alterations between children with FAS and PFAS and those who are heavily exposed but nonsyndromal have recently been reported (Cheng et al., 2017; Diwadkar et al., 2013; Li et al., 2009; Robertson et al., 2015; Sullivan et al., 2020). However, degree of PAE and severity of cognitive impairment were often not controlled in these previous studies, which could have contributed to the observed neural differences. It is important to evaluate the brain-behavioral relations between participants with FAS or PFAS and those seen in nonsyndromal HE children independently of these confounding factors.

To address these issues, the current study examined the neural mechanisms associated with atypical reading skills in two groups of adolescents (FAS/PFAS vs. HE) with similar levels of PAE and reading proficiency, in comparison with typically developing controls. Given the critical role of phonological skills in reading acquisition (Melby-Lervåg et al., 2012; Wagner & Torgesen, 1987) and the wide range of reading skills observed in adolescents with FASD, neural responses during phonological processing were examined in an fMRI study. In addition, diffusion tensor imaging (DTI) data were acquired and analyzed to characterize the structural mechanisms underlying (atypical) reading development in participants with FASD. Based on the previous findings of the PAE-associated reading deficits and brain alterations, we predicted that, compared to controls, individuals with FASD would show atypical functional responses during phonological processing and white matter disruptions in reading-related tracts. Given the previously identified associations between PAE-related facial dysmorphologies and brain structure, we expected that syndromal participants, i.e., those with FAS and PFAS, would differ in neural correlates of reading performance, compared with nonsyndromal adolescents with HE.

## 2. Methods

### 2.1. Participants

The sample consisted of 75 adolescents from the Cape Coloured (mixed ancestry) community in Cape Town, South Africa. 59 were from the Cape Town Longitudinal Cohort (S. W. Jacobson et al., 2008), for which women were initially recruited during pregnancy. To increase sample size, 16 additional adolescents were recruited from classrooms from the same public schools attended by the adolescents in the original cohort. 26 of the 75 participants were diagnosed with FAS (*n* = 10) or PFAS (*n* = 16) following a standard protocol (Hoyme et al., 2005; S. W. Jacobson et al., 2021). 30 adolescents whose mothers reported heavy drinking during pregnancy but did not meet criteria for FAS or PFAS were classified as non-syndromal heavily exposed (HE). 19 participants were controls with reading proficiency in the normal range (defined below), whose mothers had abstained from alcohol use during pregnancy. The three groups were balanced by age and sex (Table 1). All participants were right-handed, except for 4 left-handed (3 HE and 1 control) and 4 ambidextrous (1 FAS/PFAS, 1 HE and 2 controls). Approval for human research was obtained from the Wayne State University and University of Cape Town Institutional Review Boards with a reliance agreement at Boston Children’s Hospital. Written informed consent was obtained from the adolescent’s parent or guardian; written assent, from the adolescent.

**Table 1.** Sample characteristics.

|  | FAS/PFAS | HE | Control | Group effect |
| --- | --- | --- | --- | --- |
| Number | 26 | 30 | 19 | - |
| Sex (Female/Male) | 10/16 | 9/21 | 7/12 | $\chi^2 = 0.49$ |
| Age (years) | $16.8 \pm 0.7$ | $16.4 \pm 1.2$ | $16.3 \pm 1.1$ | $F_{2,72} = 1.4$ |
| Family socio-economic status | $16.7 \pm 6.6^a$ | $19.5 \pm 6.7^a$ | $24.6 \pm 7.6^b$ | $F_{2,72} = 7.3^{***}$ |
| ADHD comorbidity (Yes/No) | 11/15 | 13/17 | 8/11 | $\chi^2 = 0.01$ |
| Language (English/Afrikaans) | 8/18 | 17/13 | 17/2 | $\chi^2 = 15.4^{**}$ |
| **Alcohol consumption, smoking and drug use during pregnancy** |  |  |  |  |
| Absolute alcohol (oz)/day | $1.2 \pm 1.4$ | $0.9 \pm 1.1$ | - | $t_{54} = 0.72$ |
| Absolute alcohol (oz)/drinking day | $4.1 \pm 1.7$ | $3.9 \pm 1.9$ | - | $t_{54} = 0.43$ |
| Frequency of drinking (days/week) | $1.8 \pm 1.2$ | $1.5 \pm 1.1$ | - | $t_{54} = 1.1$ |
| Cigarettes (#/day) | $7.9 \pm 6.3^a$ | $8.1 \pm 6.7^a$ | $1.8 \pm 2.9^b$ | $F_{2,72} = 7.9^{***}$ |
| Marijuana (days/week) | $0.2 \pm 0.62$ | $0.8 \pm 1.8$ | - | $t_{54} = 1.8$ |
| **Psychometric assessment** |  |  |  |  |
| TOWRE: Phonetic Decoding Efficiency | $74.2 \pm 16.0^a$ | $81.9 \pm 18.9^a$ | $95.2 \pm 10.3^b$ | $F_{2,69} = 8.9^{***}$ |
| TOWRE: Sight Word Efficiency | $72.3 \pm 12.0^a$ | $77.3 \pm 12.7^a$ | $92.8 \pm 6.9^b$ | $F_{2,71} = 18.8^{***}$ |
| WASI IQ | $68.7 \pm 9.9^a$ | $81.3 \pm 13.4^b$ | $86.2 \pm 15.4^b$ | $F_{2,72} = 11.6^{***}$ |
| **fMRI task performance** |  |  |  |  |
| Accuracy (% correct) | $93.4\% \pm 0.09$ | $92.3\% \pm 0.11$ | $96.1\% \pm 0.04$ | $F_{2,69} = 1.14$ |

Note. Values are mean ( ± standard deviation) for continuous measures and frequencies for
hoc tests.

Note. Values are mean (± standard deviation) for continuous measures and frequencies for dichotomous measures. Superscripts indicate groups that did not differ from each other on post hoc tests.

FAS: fetal alcohol syndrome; PFAS: partial fetal alcohol syndrome; HE: non-syndromal heavily exposed; TOWRE: Test of Word Reading Efficiency; WASI: Wechsler Abbreviated Scale of Intelligence. \**p* < 0.05; \*\**p* < 0.01; \*\*\**p* < 0.001.

### 2.2. Assessment of prenatal alcohol exposure and drug use

The Cape Town Longitudinal Cohort was recruited during pregnancy between 1999-2002 (S. W. Jacobson et al., 2008) from a local antenatal clinic that serves an economically disadvantaged mixed ancestry population in which heavy drinking during pregnancy is highly prevalent (Croxford & Viljoen, 1999; May et al., 2013). Mothers were interviewed at recruitment, mid-pregnancy and 1-month postpartum about their drinking behaviors on a day-by-day basis using a timeline follow-back approach (S. W. Jacobson et al., 2002). Volume was recorded for each type of alcoholic beverage consumed each day and converted to ounces (oz) of absolute alcohol (AA) using multipliers proposed by Bowman et al. (1975, liquor = 0.4, wine = 0.12, beer = 0.05, cider = 0.06). Three summary measures were constructed: average oz AA/day, oz AA/drinking occasion, and days/week of alcohol consumption. Heavy drinking was defined as ≥14 standard drinks/week (1 oz ≈ 2 standard drinks) or binge drinking, which at that time was defined as ≥5 drinks/occasion during pregnancy. All women who reported heavy drinking during pregnancy were advised to stop or reduce their intake and were offered referral for treatment. Women <18 years of age and those with diabetes, epilepsy, or cardiac problems requiring treatment were not included in the study.

Mothers of the 16 additional adolescents were interviewed retrospectively regarding their current and pregnancy alcohol consumption using timeline follow-up interviews. All 75 mothers were administered the Structured Clinical Interview for DSM-IV Disorders (SCID-I; First et al., 1995) to diagnose lifetime alcohol abuse and/or dependence and were asked how many years they had been drinking and the largest quantity of alcohol they had consumed on a single occasion. To ensure that the assessment of PAE was comparable for both groups, data from the prospectively-recruited mothers were examined in linear regression models constructed to “predict” alcohol use during pregnancy from current drinking, retrospectively-reported drinking during pregnancy, history of alcohol abuse or dependence (yes/no), years of drinking, and most drinks on a single occasion. Multiple *R*s for AA/day, AA/occasion, and days/week of drinking were 0.68, 0.66, and 0.76, respectively. These regression models were then used to estimate the three measures of maternal alcohol use during pregnancy for the participants recruited during adolescence. All the mothers were also asked how many cigarettes they smoked/day and how many days/week they used marijuana (“dagga”), methaqualone (“mandrax”), cocaine, and any other illicit drugs during pregnancy.

### 2.3. FAS/PFAS diagnosis

In September 2005, we organized a clinic in which each child from the prospectively recruited cohort was independently examined for growth and the anomalies seen in FAS and PFAS using a standard protocol (Hoyme et al., 2005) by two expert FAS dysmorphologists (H. Eugene Hoyme, MD, and Luther K. Robinson, MD), who subsequently reached agreement regarding FASD diagnosis (see (S. W. Jacobson et al., 2021; S. W. Jacobson et al., 2008). Interobserver reliability between these dysmorphologists was substantial, including palpebral fissure length and philtrum and vermilion ratings based on the Astley and Clarren (2001) rating scales (*r*’s = 0.80, 0.84, and 0.77, respectively). Participants were re-examined by the same two dysmorphologists in a second clinic in 2009 (Suttie et al., 2013) and in clinics led by Dr. Hoyme in 2013 and 2016. Dr. Hoyme subsequently reviewed the diagnoses from all four assessments in case conferences with Drs. S. and J. Jacobson, and they assigned a final diagnosis to each participant (S. W. Jacobson et al., 2021). The 16 adolescents who were recruited retrospectively were diagnosed by Dr. Hoyme using the same protocol in 2017.

### 2.4. Attention-Deficit/Hyperactivity Disorder (ADHD) comorbidity

Due to the high comorbidity of ADHD and FASD (Kingdon, Cardoso, & McGrath, 2016), all participants were screened for ADHD using the Disruptive Behaviors Disorders Checklist (Pelham et al., 1992). Participants were assigned an ADHD classification following procedures we developed in collaboration with Joel Nigg, Ph.D., and Rafael Klorman, Ph.D., two licensed clinicians widely recognized for their expertise in ADHD research (see J. L. Jacobson et al., 2011). Symptom counts were computed separately for inattention and hyperactivity at each age by totaling how many of the nine DSM-IV behavioral criteria for each ADHD subtype were endorsed as ‘‘often’’ or ‘‘very often’’ by the mother and the child’s classroom teacher. An ADHD classification was assigned based on DSM-IV criteria if (a) 6 of 9 symptoms of inattention and/or 6 of 9 symptoms of impulsivity/hyperactivity was endorsed (‘‘pretty much’’ or ‘‘very much’’ true of child) by the mother or teacher and (b) at least 2 symptoms were endorsed by the other informant, ensuring that the behavior was observed in at least two different settings. Because 11 (42%) FAS/PFAS and 13 (43%) HE participants were diagnosed with ADHD, participants with ADHD were oversampled in the control group to match the ratio of ADHD diagnoses in the other two groups (*X*^2^ = 0.01, *p* > 0.9, see Table 1).

### 2.5. Behavioral and socioenvironmental assessments

Reading proficiency was assessed using the Test of Word Reading Efficiency (TOWRE, Torgesen et al., 1999), in which participants are asked to name real words (“sight word efficiency”) or pronounceable pseudowords (“phonemic decoding efficiency”) as quickly and accurately as possible within 45 seconds. All control participants scored in the normal range, defined as within or above one standard deviation from the mean of both tasks (i.e., standard scores > 85). General cognitive functioning was assessed using the Wechsler Abbreviated Scale of Intelligence (WASI) IQ test (Wechsler, 2011). These assessments were conducted by neuropsychology-trained Masters-level graduate research assistants, who were blind to the participants’ PAE.

It should be noted that because parents can choose between Afrikaans and English as their child’s primary language of instruction in the public schools in Cape Town, The psychometric assessments were conducted in the primary language (Afrikaans or English) of instruction used in each participant’s school. The Afrikaans versions of the reading assessments and the fMRI task were translated by a graduate research assistant, Landi Meiring, M.S., who majored in neuropsychology and linguistics, under the supervision of and in collaboration with Simone Conradie, Ph.D., a linguist on faculty at Stellenbosch University. Care was taken to match words for semantic category, linguistic difficulty, number of syllables, and word familiarity. Adjustments were made to the English items when Afrikaans items could not be matched to the existing items, and items were culturally adjusted where appropriate for a South African sample (e.g., ‘lion’ instead of ‘bear’). Moreover, to ensure that the observed effects were independent of language influences, additional analyses were performed on the participants who were tested using the English version of the assessments and fMRI tasks to see if they were comparable to the overall findings (see *Follow-up analyses based on data of English-assessed participants only*).

Socioeconomic status (SES) was assessed using the Hollingshead scale (Hollingshead, 2011), based on parental occupational status and years of education. Each of the continuous demographic, prenatal exposure, and behavioral measures were subjected to analysis of variance (ANOVA) with group as the between-subject factor. *Post-hoc* analyses (two-sample tests) were performed for measures with significant group effects. Dichotomous variables were analyzed using chi-square tests.

### 2.6. Imaging acquisition

Neuroimaging data were collected on a 3T Siemens Skyra MRI scanner with a standard Siemens 32-channel head coil at the Cape Universities Body Imaging Centre. Structural images were acquired using a multi echo MPRAGE sequence (van der Kouwe, Benner, Salat, & Fischl, 2008) with parameters of 176 slices, TR = 2530 ms, TEs = 1.67/ 3.52/ 5.37/ 7.22 ms, flip angle = 7°, field of view (FOV) = 230 mm^2^, voxel size = 1.0×1.0×1.0 mm^3^, GRAPPA 2. For fMRI data acquisition, a behavior interleaved gradient (BIG) imaging design was applied with the following parameters: TR = 6000 ms; TA = 2000 ms; TE = 30 ms; flip angle = 90°; FOV = 250 mm^2^; in-plane resolution = 3 × 3 mm^2^, slice thickness = 4 mm, slice gap = 1 mm. Diffusion data were collected in two sequences with opposite phase encoding directions (i.e., anterior–posterior and posterior–anterior). Each sequence included 5 non-diffusion-weighted volumes (b=0) and 30 diffusion-weighted volumes acquired with non-colinear gradient directions (b=1000 s/mm^2^), all at 122x122 base resolution and isotropic voxel resolution of 2.0 mm isotropic.

### 2.7. fMRI

#### 2.7.1. fMRI task procedure

A first sound matching (FSM) task was presented in a block design to examine the neural mechanisms underlying phonological processing (Raschle et al., 2012, 2013; Yu et al., 2018, 2020). During each trial, the participant saw two pictures of two common objects presented on the left and right side of the screen for 2 seconds each, accompanied by the objects’ names spoken by either a male or a female. Both pictures remained on the screen for another 2 seconds, during which the participant indicated whether the two names shared the same first sound (e.g., goat vs. gorilla) or had a different first sound by pressing the left- or right-hand button, respectively, on a key pad. In the voice-matching (VM) control task, the participant pressed the left button to indicate that the two object names were spoken by the same person (i.e., both by the male or both by the female); the right, by a different person. The order of the two tasks was counterbalanced across participants. Each trial lasted 6 seconds, equal to the length of one TR. Application of the BIG design ensured that the actual scanning time was synchronized with the response period, allowing the auditory stimuli to be presented when the scanner background noise was reduced (Gaab et al., 2007a, 2007b, 2008; Hall et al., 1999). Each task block was comprised of 4 trials, lasting 24 seconds. Each FSM run consisted of seven task blocks alternating with seven fixation blocks of the same length (24s).

#### 2.7.2. fMRI image analyses

Functional imaging data were acquired from all participants. Data were excluded for one subject due to incidental findings; one, poor image quality; and one, a technical problem. fMRI data from the remaining 72 participants (25 FAS/PFAS, 29 HE, 18 controls) were analyzed in SPM8 (http://www.fil.ion.ucl.ac.uk/spm/software/spm8), based on Matlab (Mathworks). A standard preprocessing pipeline was applied, which included 1) removal of initial volumes due to T1 equilibration effects; 2) correction for head movement (realignment); 3) normalization to the Montreal Neurological Institute (MNI) space via the high-resolution structural images using the CAT12 toolbox (Gaser & Dahnke, 2016); 4) smoothing with a Gaussian kernel with full-width at half maximum of 8 mm. Moreover, scans with excessive motion, defined as 1.5 mm (translational) and/or 2° (rotational) from the beginning scan of each run, were identified. All participants had less than 15% scans with excessive motion (mean = 0.7% ± 0.02). The three groups did not differ significantly in either the number of removed scans (*F*2,69 = 1.2, *p* = 0.3) or the head movement of remaining images (*F*2,69 = 1.1, *p* = 0.3). Preprocessed images were then submitted to a general linear model with experimental (FSM and VM) and rest conditions as regressors of interest, along with nuisance covariates for run effect and an intercept term. To minimize the potential effects of head movement, six motion parameters and binary regressors, each coding one image with excessive motion, were also included. After model estimation, neural responses for phonological processing were computed by contrasting the beta map of FSM with that of VM for each participant.

The neural correlates of phonological processing in the three groups (FAS/PFAS, HE, controls) were investigated using two analytic approaches assessing the group-level characteristics and inter-subject variability, respectively. Specifically, a one-way ANOVA model was first built with an *F* contrast to identify regions showing differences in the mean activation magnitudes between the three groups. Second, the potential differences in the brain-behavior relations between the three groups were examined in analyses of covariance (ANCOVA) for the behavioral variables “phonemic decoding efficiency” and “sight word efficiency performance”, respectively. Each ANCOVA model included three binary regressors for the three participant groups and three continuous regressors, each representing the individual performance on the phonemic decoding efficiency or sight word efficiency tests within one group. An *F* contrast derived from the continuous regressors was then computed to identify neural regions with a significant group by reading performance interaction effect, indicating differentiated neural mechanisms underlying the individual differences between the three groups. For both analyses, the significant whole-brain results were reported at a cluster-level significance of *p* < 0.05 (voxel-level *p* < 0.001), Monte-Carlo corrected for multiple comparisons.

To further explore the specific patterns of the significant group effect observed in the whole brain analyses, region-of-interest (ROI) analyses were subsequently conducted. Mean contrast estimates of FSM > VM were extracted for every participant in each identified ROI. For regions with significant ANOVA effects, two-sample *t*-tests were performed to assess the differences in activation magnitudes for each group pair (i.e., FAS/PFAS vs. HE, HE vs. controls, FAS/PFAS vs. controls). The same analyses were also performed for each group pair to examine whether the neural associations with the reading scores (phonemic decoding efficiency or sight word efficiency) were different between FAS/PFAS and HE, HE and controls, FAS/PFAS and controls in each of the ANCOVA-derived ROIs. These analyses were followed by correlation analyses relating neural activation magnitudes to the reading scores within each group in order to interpret any observed interaction effects.

### 2.8. Diffusion image analyses

In contrast to the left-hemispheric dominance that characterizes typical reading development, the current fMRI experiment revealed right-hemispheric reliance in adolescents with FAS and PFAS during phonological processing (see *fMRI results below*). This finding suggests atypical functional lateralization associated with PAE-associated reading impairments, which is consistent with the reduced brain asymmetries associated with FASD reported previously. We, therefore, set out to systematically examine structural lateralization in the key white matter tracts underlying reading development. DTI data collected from 70 participants were first corrected for susceptibility distortions, eddy current and subject movements using the FSL toolbox (https://fsl.fmrib.ox.ac.uk/fsl/fslwiki). Volumes with artifacts caused by persistent head motion (>2mm/0.5°) after correction, interlace-wise “venetian blind”, slice-wise and gradient-wise intensity inconsistencies were removed (DTIprep, Liu et al., 2010). Three adolescents (2 FAS/PFAS, 1 control) with more than 15% gradients removed were excluded, resulting in a final sample of 67 participants (23 FAS/PFAS, 27 HE, 17 controls). The three groups did not differ significantly in either the number of removed gradients (*F*2,64 = 0.8, *p* = 0.4) or head movement in the remaining images (*F*2,64 = 1.7, *p* = 0.2). The corrected DTI images were aligned to the corresponding structural images via the b0 images. They were then fitted using a linear least-squares fit, and fractional anisotropy (FA) maps were calculated for all subjects (Basser et al., 1994).

White matter tracts were identified using the Automated Fiber Quantification toolbox (Yeatman et al., 2012). The four white matter tract pairs that underlie the neural reading network were reconstructed; namely, the arcuate fasciculus (AF) that connects the inferior frontal with the middle temporal lobe, superior longitudinal fasciculus (SLF) linking the inferior parietal with the anterior frontal areas, as well as inferior longitudinal fasciculus (ILF) and inferior frontal-occipital fasciculus (IFOF) that underlie the temporo-occipital circuits with the latter extending to the anterior frontal regions (Figure 1). Whole-brain tractography was first estimated using a deterministic streamline tracking algorithm (Basser et al., 2000) with FA = 0.2/angle = 40° termination thresholds. Using two pre-defined waypoint ROIs (transformed from the MNI space to the native space via T1), eight fiber tracts of interest were then successfully delineated in all participants, except one (FAS/PFAS) for the left ILF and nine (4 FAS/PFAS and 5 HE) for the right AF. Tract refinement was subsequently performed with reference to the manually segmented fiber tract probability maps (Hua et al., 2008) and then tracts were sampled to 100 equidistant nodes. FA, as an indication of the fiber directionality, was estimated for each node, producing a detailed “tract profile” for each tract. We then computed the left-lateralization index (LI) of each node using the following formula: LI = [left FA– right FA]/[left FA + right FA] (Vandermosten et al., 2013).

**Figure 1.**
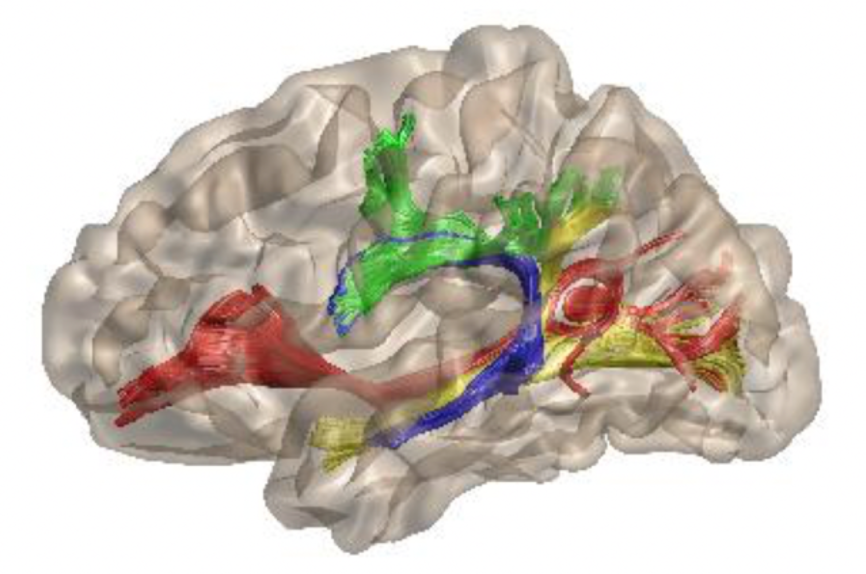
3D projections of the four white matter tracts underlying the canonical reading circuits, including the arcuate fasciculus (blue), superior longitudinal fasciculus (green), inferior longitudinal fasciculus (yellow) and inferior frontal-occipital fasciculus (red). Only left-hemispheric tracts are present for better illustration.

As in the fMRI analyses, the LI measures of each node were subjected to the ANOVA and ANCOVA models to evaluate group differences in left-lateralized magnitudes and the associations with reading performances (phonemic decoding efficiency and sight word efficiency). Significant results were reported at a cluster-level *p* < 0.05 (a node-level *p* < 0.05), family-wise error corrected for multiple comparisons (Nichols & Holmes, 2002). In ROI analyses, two-sample *t*-tests were conducted for each group pair (FAS/PFAS vs. HE, HE vs. controls, FAS/PFAS vs. controls) in white matter segments with significant ANOVA results. For ANCOVA-derived results, interaction effects were examined for each group pair, followed by correlation analyses within each group. Since the LI reflected comparisons between left- and right-hemispheric tract characteristics, FA values of hemisphere-specific tract segments were further analyzed for the white matter segments with significant results to identify the group differences in each tract.

## 3. Results

### 3.1. Sample characteristics (Table 1)

SES was higher for the control group than in both the FAS/PFAS (*t*43 = 3.7; *p* < 0.001) and HE (*t*47 = 2.5; *p* = 0.02) groups, and the latter two groups did not differ significantly from each other (*t54* = 1.6; *p* = 0.12). Maternal alcohol consumption during pregnancy was similar in the FAS/PFAS and HE groups in terms of average AA/day, AA/occasion and frequency (all *p*s > 0.2), and all of the control mothers abstained from drinking alcohol during pregnancy. Mothers of adolescents from the FAS/PFAS and HE groups smoked similar numbers of cigarettes per day (*t*54 = 0.1, *p* = 0.9), and both groups smoked more than the mothers of the control participants (FAS/PFAS vs. Control: *t*43 = 3.9, *p* < 0.001; HE vs. Control: *t*47 = 3.9, *p* < 0.001). Marijuana use during pregnancy was reported by 9 mothers (2 FAS/PFAS and 7 HE) with no significant group differences in frequency, and 4 mothers reported using illicit drugs (1 from the FAS/PFAS group used cocaine 2.6 days/week during pregnancy; 3 from the HE group used methaqualone 1.9 ± 1.7 days/week).

Reading performance scores were successfully obtained from all but one subject (FAS/PFAS) for the sight word efficiency (SWE) test and all but three (one from each group) for the phonemic decoding efficiency (PDE) test. Significant group effects were observed on both tests. *Post-hoc* analyses revealed the same pattern for both tests; while the FAS/PFAS and HE groups did not differ from each other (SWE: *t*53 = 1.5; *p* = 0.15; PDE: *t*52 = 1.6, *p* = 0.11), both groups performed more poorly than the controls (FAS/PFAS vs. Control: SWE: *t*42 = 6.6, *p* < 0.001, PDE: *t*41 = 4.9, *p* < 0.001; HE vs. Control: SWE: *t*47 = 4.9, *p* < 0.001, PDE: *t*45 = 2.7, *p* = 0.009).

The WASI IQ test also showed a significant group effect, but here the FAS/PFAS group performed more poorly than both the HE (*t*54 = 4.0, *p* < 0.001) and control (*t*43 = 4.6, *p* < 0.001) groups, and the latter two groups did not differ from each other (*t*47 = 1.2, *p* = 0.25). Finally, it should be noted that, while Afrikaans was the primary language for the FAS/PFAS group, English was the primary language for the control group (Table 1). Therefore, the replication analyses based on only the data of participants assessed in English were further conducted for the observed findings. These analyses were done to ensure that the observed results reflect neural characteristics inherent to FASD instead of being driven by language differences (see *Results 3.4. Follow-up analyses based on data of English-assessed participants only*).

### 3.2. fMRI results

#### 3.2.1. Behavioral performance

Adolescents from all three groups performed well on the phonological processing task (mean accuracies = 94% ± 0.09) with no significant differences between groups (Table 1).

#### 3.2.2. fMRI imaging results (Figure 2A and 2B)

The *F* contrast derived from the ANOVA model revealed a significant group effect on activation magnitudes among the three groups in the right precentral gyrus (RPrecentral, [42, -9, 63], k = 41). The ROI analyses showed significantly higher activation for the FAS/PFAS than for both the HE (*t*52 = 4.9, *p* < 0.001) and control groups (*t*41 = 3.0, *p* = 0.005), whereas the latter two groups did not differ from each other (*t*45 = 0.8, *p* = 0.4).

**Figure 2.**
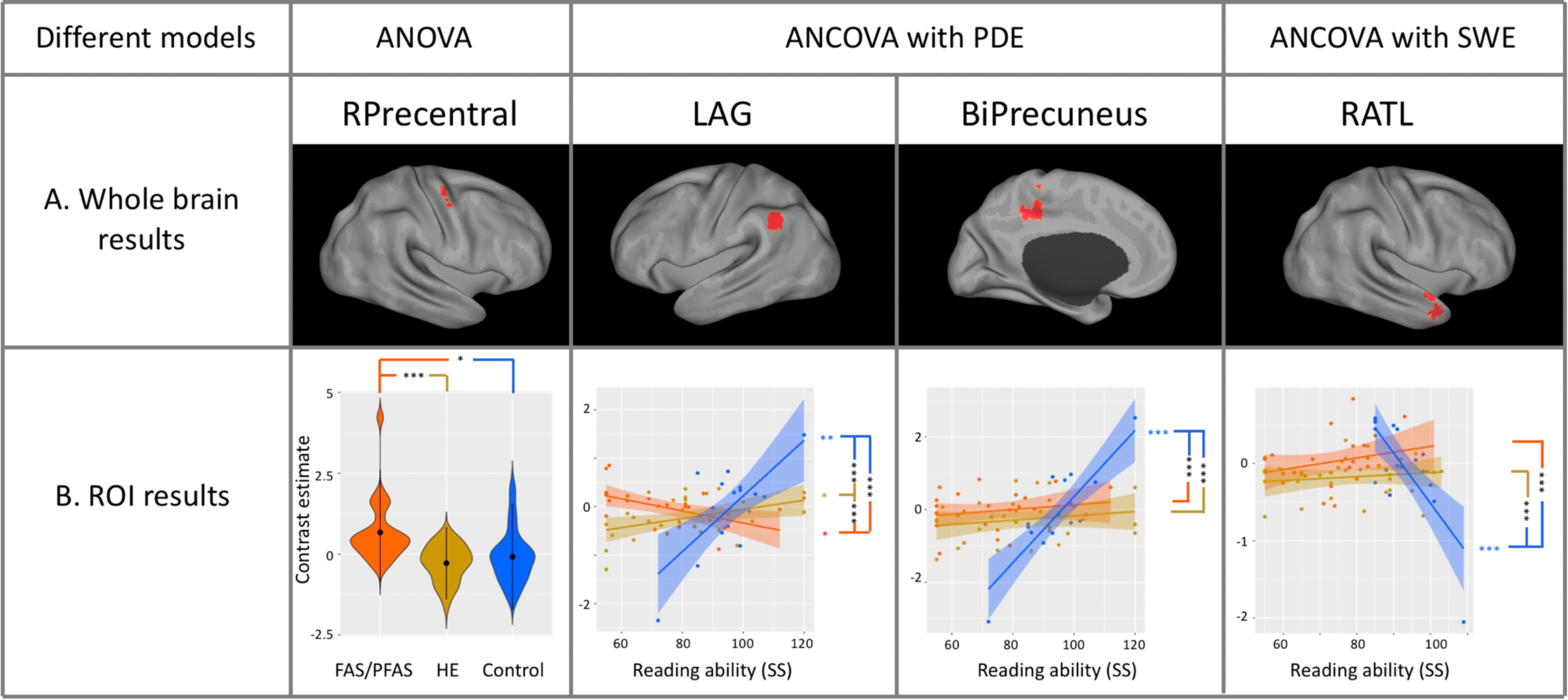
Distinctive neural responses during the phonological processing task between adolescents with full or partial fetal alcohol syndrome (FAS/PFAS, red), non-syndromal heavily exposed (HE, yellow), and controls (blue). Panel A shows the whole-brain results derived from 1) the ANOVA tests that evaluated the differences in the activation magnitudes among the three groups; 2) the ANCOVA tests that examined the group differences in the associations between the neural activation and the individual differences in reading skills, measured by the phonemic decoding efficiency (PDE) and sight word efficiency (SWE) tests, respectively. The whole-brain results were reported at a cluster-level significance of p < 0.05 (voxel-level p < 0.001), Monte-Carlo corrected for multiple comparisons. Panel B illustrates the group-specific neural characteristics in each region and the results of pairwise comparisons (FAS/PFAS vs. HE, HE vs. controls, FAS/PFAS vs. controls) based on the two-sample *t*-tests and ANCOVA for the whole brain ANOVA and ANCOVA results, respectively. Moreover, in regions derived from the ANCOVA analyses, results of correlations of neural measures with reading performance are also presented to further explain the observed group differences. * *p* < 0.05, ** *p* < 0.01, *** *p* < 0.001.

The *F* effect derived from the ANCOVAs revealed significant group differences in the neural correlates of performance on the phonemic decoding efficiency test in the left angular gyrus (LAG, [-51, -54, 36], k = 71) and bilateral precuneus cortex (BiPrecuneus, [-3, -48, 48], k = 168). ROI analyses of the LAG revealed significant interaction effects between the FAS/PFAS and control groups (*F*1,37 = 28.2, *p* < 0.001) and between the FAS/PFAS and HE groups (*F*1,48 = 13.1, *p* = 0.001), driven by the negative association between group and phonemic decoding efficiency scores in the FAS/PFAS group (*r*22 = -0.50, *p* = 0.01) in contrast to the positive associations in the HE (*r*15 = 0.70, *p* = 0.002) and control (*r*26 = 0.44, *p* = 0.019) groups (Fig. 1B). There was also a significant interaction effect between the HE and control groups (*F*1,41 = 14.4, *p* < 0.001) due to a greater positive association between neural magnitude and PDE scores in the control than the HE group. For BiPrecuneus, significant interaction effects were observed between group and phonemic decoding efficiency scores when controls were compared with the FAS/PFAS (*F*1,37 = 28.1, *p* < 0.001) and HE groups (*F*1,41 = 29.7, *p* < 0.001) due to the significantly positive correlation in the controls (*r15* = 0.84, *p* < 0.001) not seen in the FAS/PFAS (*r22* = 0.22, *p* = 0.31) and HE (*r26* = 0.17, *p* = 0.39) groups. The FAS/PFAS and HE groups did not differ in their associations of BiPrecuneus activations with the phonemic decoding efficiency scores (*F*1,48 = 0.05, *p* = 0.83).

The ANCOVAs for the sight word efficiency scores revealed a significant interaction effect in the right anterior temporal lobe (RATL, [54, 9, -30], k = 37). ROI analyses showed that both the FAS/PFAS and HE groups differed from the controls in the associations between neural activation and single word efficiency scores (FAS/PFAS: *F*1,38 = 27.9, *p* < 0.001; HE: *F*1,43 = 30.2, *p* < 0.001), due to a negative correlation in the controls (*r16* = -0.75, *p* < 0.001) not seen in either the FAS/PFAS (*r22* = 0.30, *p* = 0.16) or HE (*r27* = 0.12, *p* = 0.54) groups. No significant interaction effect was observed for the comparison between the FAS/PFAS and HE groups (*F*1,49 = 0.70, *p* = 0.41).

### 3.3. DTI results (Figure 3)

ANOVAs of the LI values for the four reading-related tracts identified significant group effects in the middle portion of the ILF (nodes 48-62). ROI analyses showed significant lower LI values for the FAS/PFAS than the HE (*t*47 = 2.0, *p* = 0.05) and control (*t*37 = 2.9, *p* = 0.006) groups, whereas the latter two groups were not different from each other (*t*42 = 1.5, *p* = 0.14). Analyses of the hemisphere-specific FA metrics showed a significant group effect in the right ILF (*F*2,64 = 3.6, *p* = 0.03) due to higher FA values in the FAS/PFAS than the HE (*t*48 = 2.4, *p* = 0.02) and control (*t*38 = 2.4, *p* = 0.02) groups; whereas the HE and control groups did not differ from each other (*t*42 = 0.24, *p* = 0.82). No significant group differences were observed for the left ILF (*F*2,63 = 0.52, *p* = 0.6).

**Figure 3.**
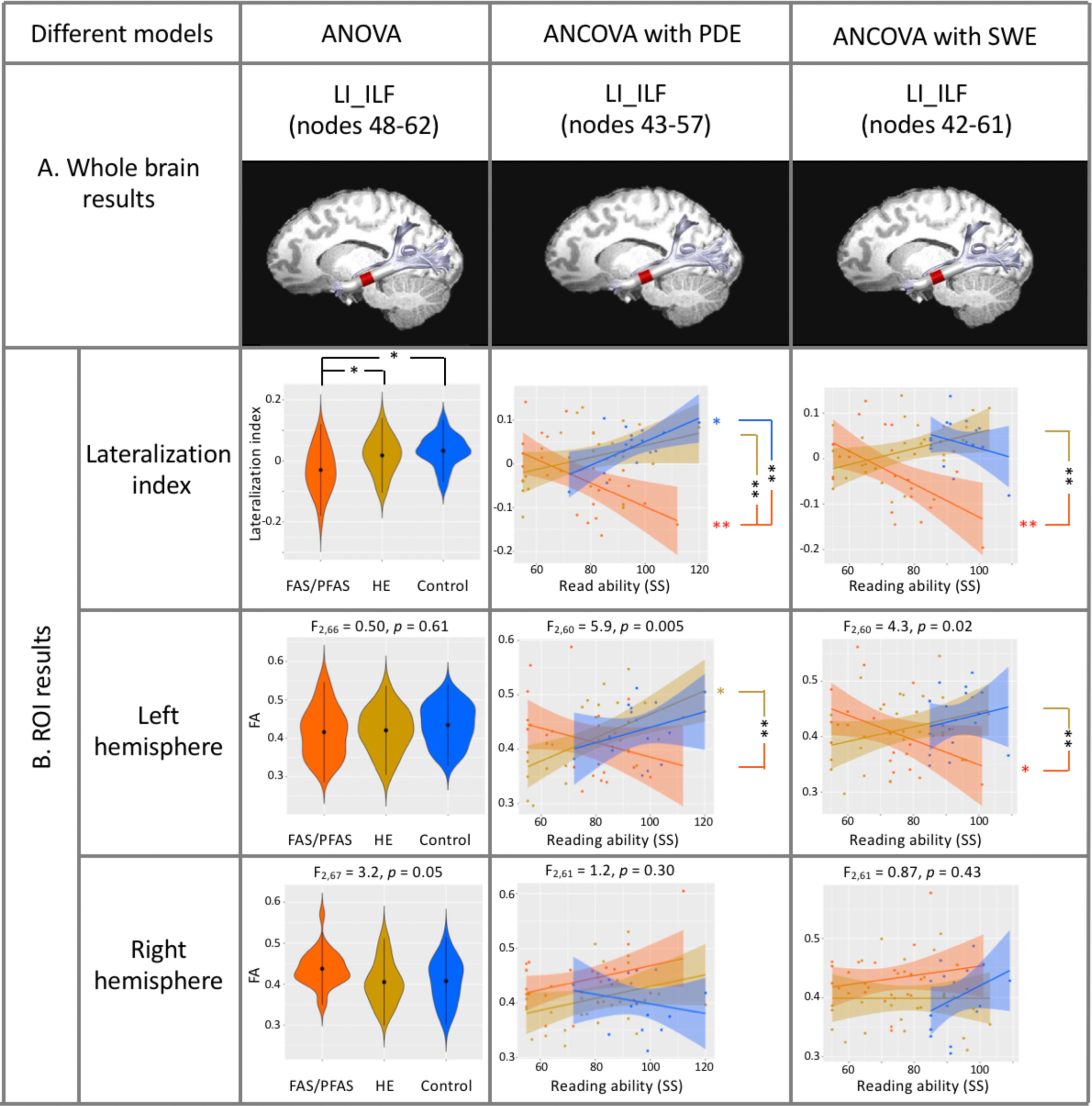
Atypical hemispheric- lateralization of the inferior longitudinal fasciculus (ILF) in adolescents with full or partial fetal alcohol syndrome (FAS/PFAS, red), as compared to those in non-syndromal heavily exposed (HE, yellow) and controls (blue). Panel A presents the specific segments of the ILF that show significant group differences in the left-lateralization index (LI), as revealed by the ANOVA tests, as well as the associations between the LI values and the reading performance of the phonemic decoding efficiency (PDE) and sight word efficiency (SWE) tests, as demonstrated by the ANCOVA tests. Significant DTI results were reported at a cluster-level *p* < 0.05 (a node-level *p* < 0.05), family-wise error corrected for multiple comparisons. Panel B illustrates the group-specific white matter characteristics, including the LI values and hemispheric-specific fractional anisotropy (FA) values in each identified segment. For the LI measures, significant between-group differences in LI values and associations between LI and reading measures were shown. For hemispheric-specific measures, both the main effects of group and significant pairwise comparisons are shown. * *p* < 0.05, ** *p* < 0.01, *** *p* < 0.001.

Similar segments of ILF showed significant interaction effects between group and reading performance on the phonemic decoding efficiency (nodes 43-57) and sight word efficiency (nodes 42-61) tests. ROI analyses revealed that the neural associations with the phonemic decoding efficiency scores differed between the FAS/PFAS and HE groups (*F*1,44 = 10.8, *p* = 0.002), and between the FAS/PFAS and control groups (*F*1,34 = 10.5, *p* = 0.003), but not between the HE and control groups (*F*1,38 = 0.90, *p* = 0.35). These interaction effects were driven by group-specific association patterns between the LI values and the phonemic decoding efficiency scores, which were positive in the control (*r*14 = 0.60, *p* = 0.015), negative in the FAS/PFAS (*r*20 = -0.55, *p* = 0.009), and nonsignificant in the HE groups (*r*24 = 0.31, *p* = 0.12). Follow-up analyses based on the FA values of the corresponding segment in each hemisphere identified a significant interaction effect in the left (*F*2,58 = 5.8, *p* = 0.005). This effect was driven by significant differences in the neural association patterns with the phonemic decoding efficiency scores between the FAS/PFAS and HE (*F*1,44 = 10.4, *p* = 0.002) groups, due to a significantly positive association in the HE (*r*24 = 0.55, *p* = 0.004) but not the FAS/PFAS (*r*20 = -0.32, *p* = 0.14) group. The other two comparisons were not significant (FAS/PFAS vs. controls: *F*1,34 = 2.71, *p* = 0.11; HE vs. controls: *F*1,38 = 0.26, *p* = 0.61), nor was the association between the FA values of the ILF and the phonemic decoding efficiency scores in the controls (*r*14 = 0.30, *p* = 0.26). Finally, no significant interactions were seen in the right hemisphere (*F*2,59 = 1.3, *p* = 0.28).

For the ILF segment showing significant interaction effects with the sight word efficiency scores, ROI analyses identified significant differences in the neural associations with task performance between the FAS/PFAS and HE groups (*F*1,45 = 12.5, *p* = 0.001), driven by a significantly negative association in the FAS/PFAS (*r*20 = -0.56, *p* = 0.007), but not HE (*r*25 = 0.33, *p* = 0.09) group. No significant interaction effects were found in the other two comparisons (FAS/PFAS vs. controls: *F*1,35 = 0.47, *p* = 0.50; HE vs. controls: *F*1,40 = 2.6, *p* = 0.12), and the correlation in the control group was not significant (*r*15 = -0.28, *p* = 0.29). Further analyses based on the hemisphere-specific FA values revealed the same pattern in the left ILF (*F*2,60 = 3.9, *p* = 0.025), where a significant interaction effect was observed only when FAS/PFAS and HE were compared (*F*1,45 = 7.0, *p* = 0.011), but not in the other two group comparisons (FAS/PFAS vs. controls: *F*1,35 = 2.4, *p* = 0.13; HE vs. controls: *F*1,40 = 0.001, *p* = 0.98). This pattern of effects was due to a significant negative association in the FAS/PFAS (*r*20 = -0.42, *p* = 0.05) not seen in the HE (*r*25 = 0.31, *p* = 0.11) or control (*r*15 = 0.19, *p* = 0.46) groups. Finally, no significant interaction effect was evident in the right ILF (*F*2,61 = 0.88, *p* = 0.42).

### 3.4. Follow-up analyses based on data of English-assessed participants only

Afrikaans was the primary language in the FAS/PFAS group; English was the primary language for the controls. We, therefore, performed follow-up analyses of the fMRI data from the 40 adolescents (8 FAS/PFAS, 16HE, 16 controls) who were tested in English to ensure the observed group differences were not caused by language differences. The same group effect was seen in the RPrecentral ROI (*F*2,37 = 3.2, *p* = 0.05) and the same interaction effects between group and phonemic decoding efficiency score were seen in both the LAG (*F*2,32 = 4.5, *p* = 0.02) and BiPrecuneus (*F*2,32 = 7.1, *p* = 0.003). Similarly, the same interaction between group and sight word efficiency score was seen in the RATL (*F*2,34 = 4.6, *p* = 0.02).

Re-analyses of the DTI data from the 37 participants tested in English (7 FAS/PFAS, 15 HE, 15 controls) also yielded the same results as those based on the whole sample, including a significant group effect on the mean LI values of the middle segment (nodes 48-62) of the ILF (*F*2,34 = 15.8, *p* < 0.001). Significant interactions between group and phonemic decoding efficiency score were seen in the nodes 43-57 of the ILF (*F*2,29 = 4.5, *p* = 0.02) and between group and sight word efficiency score in the nodes 42-61 of the ILF (*F*2,31 = 4.9, *p* = 0.01).

## 4. Discussion

This study is the first to identify distinctive neural mechanisms associated with atypical reading skills and the associated sub-components in adolescents with FAS/PFAS and nonsyndromal HE adolescents when compared to nonexposed controls. Specifically, using a phonological processing task that assessed sub-components of reading acquisition, we observed significantly greater activation in the right precentral gyrus (RPrecentral) in adolescents in the FAS/PFAS group than both the HE and control groups. Moreover, the correlations of neural activation magnitudes during phonological processing with reading performance measured behaviorally outside the scanner were also different among the three groups. More specifically, while correlations of better performance on the phonetic decoding test with increased activation in the left angular gyrus (LAG) and the bilateral precuneus (BiPrecuneus) were observed in the controls, these associations were weaker for the HE group and in the opposite direction, i.e., increased LAG activation with lower phonetic decoding scores, in adolescents with FAS/PFAS. Finally, the correlation of better performance in the sight word efficiency test with decreased activation in the right anterior temporal lobe (RATL) among the controls was not observed in the FAS/PFAS and HE groups.

Furthermore, DTI analyses of the white matter organization relevant for reading development yielded an atypical lateralization pattern in the FAS/PFAS group. Less left-lateralization was observed in the inferior longitudinal fasciculus (ILF) in the FAS/PFAS group compared with both the HE and controls groups, an effect driven by FAS/PFAS-specific increases in FA values in the right ILF. Whereas higher FA in the left ILF was associated with better performance on both the phonemic decoding and sight word efficiency tests in the HE and control groups, it was associated with poorer performance on both reading tests in the FAS/PFAS group. No group differences were seen in the relation of FA in the right ILF to performance on either reading task. Importantly, our finding showing that the same effects were observed in the subset of participants whose primary language was English demonstrates that the observed group differences are not attributable to between-group differences in the proportion of adolescents whose primary language was Afrikaans or English. Thus, these data support our hypotheses of dissociable neural mechanisms underlying atypical reading development in adolescents with FAS and PFAS that are not seen in individuals with PAE who lack the distinctive fetal alcohol-related facial dysmorphology.

Two analytic approaches were utilized in this study to characterize different yet complementary aspects of the neural substrates underlying reading development among adolescents with FASD. The ANOVA models were applied to identify regions showing significant differences in the *group-mean* activation magnitudes among the FAS/PFAS, HE, and control groups. This approach has often been applied in previous studies on FASD (e.g., Lindinger et al., 2021; Meintjes et al., 2010; Nguyen et al., 2017) or other disabilities (e.g., developmental dyslexia, (Martin et al., 2016), with the aim of identifying the neuropathology associated with a clinical diagnosis. However, this approach does not consider *inter-subject variability* that might be meaningful in revealing etiological mechanisms. This issue was considered in our second set of analyses, which examined group differences in the correlations of neural activation magnitude or white matter organization with behavioral performance on two reading tests using the ANCOVA tests (see similar analyses in Gautam et al., 2015; Sowell, et al., 2008). These analyses examined brain-behavior relations across individuals within each of the groups, revealing the functional significance of the PAE-associated neural alterations (Lebel et al., 2011). The application of analytic methods that consider both group-level properties and individual differences provides a more comprehensive delineation of the neural anomalies in individuals with FASD exhibiting a continuum of reading impairments.

Compared to the left-lateralized neural network underlying phonological processing in typically developing individuals, the adolescents with FASD seem to recruit a more wide-spread cortical network, involving the right hemisphere. In the control group, we saw the expected positive associations between phonemic decoding scores and levels of neural activation in the LAG and BiPrecuneus, brain regions underlying phonological processing (Bitan et al., 2007; Church et al., 2008; Yu et al., 2018). Our findings that these associations were altered in adolescents with FASD are consistent with previous reports of PAE-induced structural alterations in these regions (De Guio et al., 2014; Sowell et al., 2001, 2002), suggesting high vulnerability of the left posterior temporoparietal cortex to PAE insult.

In the HE group, level of activation in the LAG was positively correlated with phonemic decoding efficiency, but the correlation was weaker than among the controls, suggesting less efficient functioning of underlying neural mechanisms, which likely contributed to their observed reading impairments. By contrast, a different neural activation pattern was seen in the participants with FAS or PFAS, who showed greater activation in the right precentral gyrus than the other two groups, suggesting increased reliance on right frontal areas. Previous studies have reported greater activation of right frontal regions in individuals with FASD during verbal learning than in controls (Sowell et al., 2007) and that verbal skills were positively correlated with cortical thickness in right frontal regions in FASD (Sowell et al., 2008), suggesting greater reliance on these regions in language skills. The current findings suggest that this verbal-related involvement of the right frontal cortex may also play a role in a critical reading-related function, i.e., phonological processing. The rightward activation was observed only in the syndromal alcohol-exposed individuals, who exhibited facial dysmorphology, implying an atypical lateralization pattern specific to this group. Consistent with this interpretation, in FASD only children with FAS have shown less left-lateralized dichotic listening asymmetry (Domellöf et al., 2009) and greater bilateral frontal activation during classical eyeblink conditioning (Cheng et al., 2017). Thus, different functional mechanisms underlying phonological processing were observed among individuals with and without facial dysmorphology; whereas a left-lateralized neural network was observed in the HE group (albeit weaker than in controls), the FAS/PFAS group relied on additional neural resources in the right hemisphere to support their reading performance.

The atypical cortical asymmetry in reading among the adolescents with FAS and PFAS is also seen in the rightward lateralization of the inferior longitudinal fasciculus (ILF) specifically observed in that group. The left ILF provides the ventral reading pathway from the temporo-occipital cortex to the anterior temporal lobe that has been associated with reading abilities among typically developing children (Vanderauwera et al., 2018; Wang et al., 2016; Yeatman et al., 2012). The ILF has previously been implicated in white matter alterations associated with PAE (Lebel et al., 2008; Sowell et al., 2008) and has been shown to have atypical developmental trajectories in individuals with FASD compared to controls (Treit et al., 2017). The current study extends these PAE-associated alterations to the lateralization characteristics of the ILF (the middle segment) specifically in the FAS/PFAS group (compared to both the HE and controls), driven by atypically higher FA values in the right ILF. It is important to note that our results are not necessarily in contrast with the previous studies that adopted a voxel-based approach and identified FA reductions in regions on the bilateral ILF pathways not overlapping with the current identified location (Fan et al., 2016; Sowell et al., 2008). Rather, our findings provide additional information about the alcohol teratogenic effects on the coordination of bilateral brain characteristics important for reading development. Specifically, these PAE-related alterations in the leftward lateralization of ILF may disrupt the development of white matter mechanisms underlying typical reading development. Accordingly, by contrast to the positive correlations between reading performance (both phonemic decoding and sight word efficiency scores) and the FA values in the left ILF in the HE and control groups, the FAS/PFAS group exhibited a negative association, suggesting atypical involvement of the ILF in reading. Overall, the white matter patterns observed in the adolescents with FAS/PFAS align with their functional characteristics during phonological processing, collectively suggesting the development of alternative brain mechanisms in the right hemisphere to compensate (albeit not entirely successfully) for PAE-associated neural alterations in the left-hemispheric reading network.

Overall, combining the fMRI and DTI findings, the current study revealed dissociable neural correlates of atypical reading development between prenatally alcohol-exposed adolescents with FAS or PFAS and those without the characteristic pattern of facial anomalies (i.e., the HE group), suggesting that distinctive brain mechanisms are employed in different subtypes of FASD. While early neuroimaging studies often combined all individuals with FASD into one group due to limited sample size (see Lebel et al., 2011; Wozniak & Muetzel, 2011 for reviews), recent investigations examining different FASD subtypes have found differences in neural activation patterns or/and structural characteristics (e.g., Astley et al., 2009; Cheng et al., 2017; Diwadkar et al., 2013). The current study is the first to provide evidence of FASD subtype-specific neural mechanisms related to reading. Importantly, since individuals in our FAS/PFAS and HE groups were similar in their levels of PAE and their reading abilities, the observed group differences cannot be attributed to different levels of exposure but instead provide compelling evidence supporting differentiated alterations in cortical engagement for reading function in adolescents with and without FAS dysmorphology. Moreover, our findings are consistent with studies that have linked more severe facial dysmorphology, such as higher lip/philtrum scores and reduced palpebral fissure length, to greater brain structural alterations (Lebel et al., 2012; Roussotte et al., 2011; Yang et al., 2012), including increased cortical thickness in the right frontal area (Yang et al., 2012).

## 5. Limitations and future directions

This study identified dissociable brain structural and functional alterations associated with reading impairments in prenatally alcohol-exposed adolescents with and without fetal alcohol-related facial dysmorphology. One potential confounding influence is our sample’s high rate of comorbidity (∼40%) of FASD with ADHD, however, it is similar to that reported in other studies (e.g., Kingdon et al., 2016). Although ADHD was oversampled in the control group to match the clinical groups, the potential influences of ADHD might have confounded group contrasts due to the unknown neural mechanisms underlying FASD and ADHD comorbidity. Future studies recruiting separate groups with single and co-morbid diagnoses are warranted to directly examine the neuropathology associated with FASD and ADHD comorbidity (see a similar study for dyslexia and ADHD comorbidity in Langer et al., 2019). Second, the participants with FASD in our study exhibited variable reading skills, with about half of the adolescents (13 FAS/PFAS and 12 HE) able to read fewer than 30 words in either reading task. Therefore, a phonological processing task was used in the current fMRI experiment. This task enabled characterization of the neural correlates of the foundational literacy skills important for reading acquisition and ensured optimal task performance during scanning. Nevertheless, further investigation with a reading comprehension task is warranted, in order to examine the neural etiologies underlying reading comprehension associated with prenatal alcohol insults and whether distinctive neural correlates are seen in different fetal alcohol-related disorders. Finally, to address potential confounds related to the primary language spoken we conducted additional analyses using data from a subset of participants with the same primary language (English) to confirm that the observed effects were independent of the language of instruction used in school. However, future studies are needed to determine whether different mechanisms underlying reading impairments are seen among individuals with different primary languages, which may have important implications for developing treatment plans more suitable for alcohol-exposed individuals with different language backgrounds.

## 5. Conclusions

In summary, utilizing both fMRI and DTI, this study is the first to characterize brain functional and structural mechanisms underlying atypical reading development and its sub- components in adolescents with prenatal alcohol insults. Specifically, the HE group showed brain-behavioral associations and left lateralization that were similar to, albeit somewhat weaker than those seen in the control group, whereas the FAS/PFAS group exhibited greater reliance on right frontal regions. Since the FAS/PFAS and HE groups in this sample were similar in terms of levels of prenatal alcohol exposure and proficiency in basic reading skills (phonemic decoding and sight word reading), these observed differences presumably reflect distinctive neural mechanisms engaged by different FASD subtypes during atypical reading and phonological processing. Given the essential role of reading proficiency for academic, vocational, economic outcomes and societal functioning generally, it is important to further understand the etiology of reading difficulties in individuals with FASD. Future studies with larger samples from socioeconomically and culturally different populations are warranted to better understand the neural etiologies of reading impairment in FASD, which might have the potential to inform the design of customized treatment strategies for PAE-exposed reading deficits.

## Data availability statement

The data sets generated during this study are available at https://osf.io/re7s9/.

## Acknowledgments

This study was supported by grants from the NIH/National Institute on Alcohol Abuse and Alcoholism RO1-AA09524 (S.J.) and R01-AA023503 (S.J. and N.G.) and the Lycaki-Young Fund from the State of Michigan (S.J. and J.J.) The authors acknowledge the assistance of Neil Dodge and Jia Fan with data collection and analyses and thank Simone Conradie, PhD, and Landi Meiring, MS, for their extensive work on the translation of the reading test into Afrikaans. The authors also express our gratitude to the families for their participation in this longitudinal study.

## References

1. Adnams, C. M., Sorour, P., Kalberg, W. O., Kodituwakku, P., Perold, M. D., Kotze, A., … May, P. A. (2007). Language and literacy outcomes from a pilot intervention study for children with fetal alcohol spectrum disorders in South Africa. Alcohol, 41(6), 403–414.

2. Astley, S. J., Aylward, E. H., Olson, H. C., Kerns, K., Brooks, A., Coggins, T. E., … Jirikowic, T. (2009). Magnetic resonance imaging outcomes from a comprehensive magnetic resonance study of children with fetal alcohol spectrum disorders. Alcoholism: Clinical and Experimental Research, 33(10), 1671–1689.

3. Astley, S. J., & Clarren, S. K. (2001). Measuring the facial phenotype of individuals with prenatal alcohol exposure: correlations with brain dysfunction. Alcohol and Alcoholism, 36(2), 147–159.

4. Basser, P. J., Mattiello, J., & LeBihan, D. (1994). MR diffusion tensor spectroscopy and imaging. Biophysical Journal, 66(1), 259–267.

5. Basser, P. J., Pajevic, S., Pierpaoli, C., Duda, J., & Aldroubi, A. (2000). In vivo fiber tractography using DT-MRI data. Magnetic Resonance in Medicine, 44(4), 625–632.

6. Ben-Shachar, M. S., Shmueli, M., Jacobson, S. W., Meintjes, E. M., Molteno, C. D., Jacobson, J. L., & Berger, A. (2020). Prenatal Alcohol Exposure Alters Error Detection During Simple Arithmetic Processing: An Electroencephalography Study. Alcoholism: Clinical and Experimental Research, 44(1), 114–124.

7. Berger, A., Shmueli, M., Lisson, S., Ben-Shachar, M. S., Lindinger, N. M., Lewis, C. E., … Jacobson, J. L. (2019). Deficits in arithmetic error detection in infants with prenatal alcohol exposure: An ERP study. Developmental Cognitive Neuroscience, 40, 100722.

8. Bitan, T., Cheon, J., Lu, D., Burman, D. D., Gitelman, D. R., Mesulam, M.-M., & Booth, J. R. (2007). Developmental changes in activation and effective connectivity in phonological processing. NeuroImage, 38(3), 564–575.

9. Bonhage, C. E., Mueller, J. L., Friederici, A. D., & Fiebach, C. J. (2015). Combined eye tracking and fMRI reveals neural basis of linguistic predictions during sentence comprehension. Cortex, 68, 33–47.

10. Bowman, R. S., Stein, L. I., & Newton, J. R. (1975). Measurement and interpretation of drinking behavior. I. On measuring patterns of alcohol consumption. II. Relationships between drinking behavior and social adjustment in a sample of problem drinkers. Journal of Studies on Alcohol, 36(9), 1154–1172.

11. Burden, M. J., Jacobson, S. W., Sokol, R. J., & Jacobson, J. L. (2005). Effects of prenatal alcohol exposure on attention and working memory at 7.5 years of age. Alcoholism: Clinical and Experimental Research, 29(3), 443–452.

12. Cattinelli, I., Borghese, N. A., Gallucci, M., & Paulesu, E. (2013). Reading the reading brain: a new meta-analysis of functional imaging data on reading. Journal of Neurolinguistics, 26(1), 214–238.

13. Cheng, D. T., Meintjes, E. M., Stanton, M. E., Dodge, N. C., Pienaar, M., Warton, C. M., … Jacobson, J. L. (2017). Functional MRI of human eyeblink classical conditioning in children with fetal alcohol spectrum disorders. Cerebral Cortex, 27(7), 3752–3767.

14. Chiodo, L. M., Jacobson, S. W., & Jacobson, J. L. (2004). Neurodevelopmental effects of postnatal lead exposure at very low levels. Neurotoxicology and Teratology, 26(3), 359–371.

15. Church, J. A., Coalson, R. S., Lugar, H. M., Petersen, S. E., & Schlaggar, B. L. (2008). A developmental fMRI study of reading and repetition reveals changes in phonological and visual mechanisms over age. Cerebral Cortex, 18(9), 2054–2065.

16. Cohen, L., Dehaene, S., Naccache, L., Lehéricy, S., Dehaene-Lambertz, G., Hénaff, M.-A., & Michel, F. (2000). The visual word form area: spatial and temporal characterization of an initial stage of reading in normal subjects and posterior split-brain patients. Brain, 123(2), 291–307.

17. Coles, C. D., Brown, R. T., Smith, I. E., Platzman, K. A., Erickson, S., & Falek, A. (1991). Effects of prenatal alcohol exposure at school age. I. Physical and cognitive development. Neurotoxicology and Teratology, 13(4), 357–367.

18. Croxford, J., & Viljoen, D. (1999). Alcohol consumption by pregnant women in the Western Cape. South African Medical Journal, 89(9).

19. De Guio, F., Mangin, J. F., Rivière, D., Perrot, M., Molteno, C. D., Jacobson, S. W., … Jacobson, J. L. (2014). A study of cortical morphology in children with fetal alcohol spectrum disorders. Human Brain Mapping, 35(5), 2285–2296.

20. Diwadkar, V. A., Meintjes, E. M., Goradia, D., Dodge, N. C., Warton, C., Molteno, C. D., … Jacobson, J. L. (2013). Differences in cortico-striatal-cerebellar activation during working memory in syndromal and nonsyndromal children with prenatal alcohol exposure. Human Brain Mapping, 34(8), 1931–1945.

21. Domellöf, E., Rönnqvist, L., Titran, M., Esseily, R., & Fagard, J. (2009). Atypical functional lateralization in children with fetal alcohol syndrome. Developmental Psychobiology: The Journal of the International Society for Developmental Psychobiology, 51(8), 696–705.

22. Donald, K. A., Roos, A., Fouche, J.-P., Koen, N., Howells, F. M., Woods, R. P., … Stein, D. J. (2015). A study of the effects of prenatal alcohol exposure on white matter microstructural integrity at birth. Acta Neuropsychiatrica, 27(4), 197–205.

23. Fan, J., Jacobson, S. W., Taylor, P. A., Molteno, C. D., Dodge, N. C., Stanton, M. E., … Meintjes, E. M. (2016). White matter deficits mediate effects of prenatal alcohol exposure on cognitive development in childhood. Human Brain Mapping, 37(8), 2943–2958.

24. Fiez, J. A., Raife, E. A., Balota, D. A., Schwarz, J. P., Raichle, M. E., & Petersen, S. E. (1996). A positron emission tomography study of the short-term maintenance of verbal information. Journal of Neuroscience, 16(2), 808–822.

25. Gaab, N., Gabrieli, J. D., & Glover, G. H. (2007a). Assessing the influence of scanner background noise on auditory processing. I. An fMRI study comparing three experimental designs with varying degrees of scanner noise. Human Brain Mapping, 28(8), 703–720.

26. Gaab, N., Gabrieli, J. D., & Glover, G. H. (2007b). Assessing the influence of scanner background noise on auditory processing. II. An fMRI study comparing auditory processing in the absence and presence of recorded scanner noise using a sparse design. Human Brain Mapping, 28(8), 721–732.

27. Gaab, N., Gabrieli, J. D., & Glover, G. H. (2008). Resting in peace or noise: Scanner background noise suppresses default-mode network. Human Brain Mapping, 29(7), 858–867.

28. Gaser, C., & Dahnke, R. (2016). CAT-a computational anatomy toolbox for the analysis of structural MRI data. Human Brain Mapping, 2016, 336–348.

29. Gautam, P., Lebel, C., Narr, K. L., Mattson, S. N., May, P. A., Adnams, C. M., … Sowell, E. R. (2015). Volume changes and brain-behavior relationships in white matter and subcortical gray matter in children with prenatal alcohol exposure. Human Brain Mapping, 36(6), 2318–2329.

30. Glass, L., Graham, D. M., Akshoomoff, N., & Mattson, S. N. (2015). Cognitive factors contributing to spelling performance in children with prenatal alcohol exposure. Neuropsychology, 29(6), 817.

31. Glass, L., Moore, E. M., Akshoomoff, N., Jones, K. L., Riley, E. P., & Mattson, S. N. (2017). Academic difficulties in children with prenatal alcohol exposure: presence, profile, and neural correlates. Alcoholism: Clinical and Experimental Research, 41(5), 1024–1034.

32. Goldschmidt, L., Richardson, G. A., Stoffer, D. S., Geva, D., & Day, N. L. (1996). Prenatal alcohol exposure and academic achievement at age six: a nonlinear fit. Alcoholism: Clinical and Experimental Research, 20(4), 763–770.

33. Hall, D. A., Haggard, M. P., Akeroyd, M. A., Palmer, A. R., Summerfield, A. Q., Elliott, M. R., … Bowtell, R. W. (1999). “Sparse” temporal sampling in auditory fMRI. Human Brain Mapping, 7(3), 213–223.

34. Hollingshead, A. B. (2011). Four factor index of social status. New Haven, CT: Yale University Press.

35. Howell, K. K., Lynch, M. E., Platzman, K. A., Smith, G. H., & Coles, C. D. (2006). Prenatal alcohol exposure and ability, academic achievement, and school functioning in adolescence: a longitudinal follow-up. Journal of Pediatric Psychology, 31(1), 116–126.

36. Hoyme, H. E., Kalberg, W. O., Elliott, A. J., Blankenship, J., Buckley, D., Marais, A.-S., … Abdul-Rahman, O. (2016). Updated clinical guidelines for diagnosing fetal alcohol spectrum disorders. Pediatrics, 138(2).

37. Hoyme, H. E., May, P. A., Kalberg, W. O., Kodituwakku, P., Gossage, J. P., Trujillo, P. M., … Khaole, N. (2005). A practical clinical approach to diagnosis of fetal alcohol spectrum disorders: clarification of the 1996 institute of medicine criteria. Pediatrics, 115(1), 39–47.

38. Hua, K., Zhang, J., Wakana, S., Jiang, H., Li, X., Reich, D. S., … Mori, S. (2008). Tract probability maps in stereotaxic spaces: analyses of white matter anatomy and tract-specific quantification. NeuroImage, 39(1), 336–347.

39. Jacobson, J. L., Dodge, N. C., Burden, M. J., Klorman, R., & Jacobson, S. W. (2011). Number processing in adolescents with prenatal alcohol exposure and ADHD: differences in the neurobehavioral phenotype. Alcoholism: Clinical and Experimental Research, 35(3), 431–442.

40. Jacobson, S. W., Chiodo, L. M., Sokol, R. J., & Jacobson, J. L. (2002). Validity of maternal report of prenatal alcohol, cocaine, and smoking in relation to neurobehavioral outcome. Pediatrics, 109(5), 815–825.

41. Jacobson, S. W., Hoyme, H. E., Carter, R. C., Dodge, N. C., Molteno, C. D., Meintjes, E. M., & Jacobson, J. L. (2021). Evolution of the physical phenotype of Fetal Alcohol Spectrum Disorders from childhood through adolescence. Alcoholism: Clinical and Experimental Research, 45(2), 395–408.

42. Jacobson, S. W., Jacobson, J. L., Sokol, R. J., Chiodo, L. M., & Corobana, R. (2004). Maternal age, alcohol abuse history, and quality of parenting as moderators of the effects of prenatal alcohol exposure on 7.5-year intellectual function. Alcoholism: Clinical and Experimental Research, 28(11), 1732–1745.

43. Jacobson, S. W., Stanton, M. E., Molteno, C. D., Burden, M. J., Fuller, D. S., Hoyme, H. E., … Jacobson, J. L. (2008). Impaired eyeblink conditioning in children with fetal alcohol syndrome. Alcoholism: Clinical and Experimental Research, 32(2), 365–372.

44. Kerns, K. A., Don, A., Mateer, C. A., & Streissguth, A. P. (1997). Cognitive deficits in nonretarded adults with fetal alcohol syndrome. Journal of Learning Disabilities, 30(6), 685–693.

45. Kingdon, D., Cardoso, C., & McGrath, J. J. (2016). Research Review: Executive function deficits in fetal alcohol spectrum disorders and attention-deficit/hyperactivity disorder–a meta-analysis. Journal of Child Psychology and Psychiatry, 57(2), 116–131.

46. Kopera-Frye, K., Dehaene, S., & Streissguth, A. P. (1996). Impairments of number processing induced by prenatal alcohol exposure. Neuropsychologia, 34(12), 1187–1196.

47. Korkman, M., Kettunen, S., & Autti-Rämö, I. (2003). Neurocognitive impairment in early adolescence following prenatal alcohol exposure of varying duration. Child Neuropsychology, 9(2), 117–128.

48. Langer, N., Benjamin, C., Becker, B. L., & Gaab, N. (2019). Comorbidity of reading disabilities and ADHD: Structural and functional brain characteristics. Human Brain Mapping, 40(9), 2677–2698.

49. Lebel, C., Mattson, S. N., Riley, E. P., Jones, K. L., Adnams, C. M., May, P. A., … Kan, E. (2012). A longitudinal study of the long-term consequences of drinking during pregnancy: heavy in utero alcohol exposure disrupts the normal processes of brain development. Journal of Neuroscience, 32(44), 15243–15251.

50. Lebel, C., Rasmussen, C., Wyper, K., Walker, L., Andrew, G., Yager, J., & Beaulieu, C. (2008). Brain diffusion abnormalities in children with fetal alcohol spectrum disorder. Alcoholism: Clinical and Experimental Research, 32(10), 1732–1740.

51. Lebel, C., Roussotte, F., & Sowell, E. R. (2011). Imaging the impact of prenatal alcohol exposure on the structure of the developing human brain. Neuropsychology review, 21(2), 102–118.

52. Li, L., Coles, C. D., Lynch, M. E., & Hu, X. (2009). Voxelwise and skeleton-based region of interest analysis of fetal alcohol syndrome and fetal alcohol spectrum disorders in young adults. Human Brain Mapping, 30(10), 3265–3274.

53. Lindinger, N., Jacobson, J., Molteno, C., Meintjes, E., Gaab, N., & Jacobson, S. (2018). The role of phonological processing, processing speed, and linguistic proficiency in reading impairment in adolescents with FASD. Alcoholism: Clinical and Experimental Research, 42, 44A.

54. Lindinger, N. M., Jacobson, J. L., Warton, C. M., Malcolm-Smith, S., Molteno, C. D., Dodge, N. C., … Jacobson, S. W. (2021). Fetal Alcohol Exposure Alters BOLD Activation Patterns in Brain Regions Mediating the Interpretation of Facial Affect. Alcoholism: Clinical and Experimental Research, 45(1), 140–152.

55. Liu, Z., Wang, Y., Gerig, G., Gouttard, S., Tao, R., Fletcher, T., & Styner, M. (2010). Quality control of diffusion weighted images. Paper presented at the Medical Imaging 2010: Advanced PACS-based Imaging Informatics and Therapeutic Applications.

56. Malins, J. G., Gumkowski, N., Buis, B., Molfese, P., Rueckl, J. G., Frost, S. J., … Mencl, W. E. (2016). Dough, tough, cough, rough: A “fast” fMRI localizer of component processes in reading. Neuropsychologia, 91, 394–406.

57. Martin, A., Kronbichler, M., & Richlan, F. (2016). Dyslexic brain activation abnormalities in deep and shallow orthographies: A meta-analysis of 28 functional neuroimaging studies. Human Brain Mapping, 37(7), 2676–2699.

58. Mattson, S. N., Bernes, G. A., & Doyle, L. R. (2019). Fetal alcohol spectrum disorders: a review of the neurobehavioral deficits associated with prenatal alcohol exposure. Alcoholism: Clinical and Experimental Research, 43(6), 1046–1062.

59. May, P. A., Blankenship, J., Marais, A.-S., Gossage, J. P., Kalberg, W. O., Joubert, B., … Hasken, J. (2013). Maternal alcohol consumption producing fetal alcohol spectrum disorders (FASD): quantity, frequency, and timing of drinking. Drug and Alcohol Dependence, 133(2), 502–512.

60. May, P. A., Chambers, C. D., Kalberg, W. O., Zellner, J., Feldman, H., Buckley, D., … Honerkamp-Smith, G. (2018). Prevalence of fetal alcohol spectrum disorders in 4 US communities. JAMA, 319(5), 474–482.

61. May, P. A., Marais, A.-S., De Vries, M. M., Buckley, D., Kalberg, W. O., Hasken, J. M., … Manning, M. A. (2021). The prevalence, child characteristics, and maternal risk factors for the continuum of fetal alcohol spectrum disorders: A sixth population-based study in the same South African community. Drug and Alcohol Dependence, 218, 108408.

62. McCandliss, B. D., & Noble, K. G. (2003). The development of reading impairment: a cognitive neuroscience model. Mental Retardation and Developmental Disabilities Research Reviews, 9(3), 196–205.

63. McLennan, J. D. (2015). Misattributions and potential consequences: The case of child mental health problems and fetal alcohol spectrum disorders. The Canadian Journal of Psychiatry, 60(12), 587–590.

64. Meintjes, E. M., Jacobson, J. L., Molteno, C. D., Gatenby, J. C., Warton, C., Cannistraci, C. J., … Gore, J. C. (2010). An FMRI study of number processing in children with fetal alcohol syndrome. Alcoholism: Clinical and Experimental Research, 34(8), 1450–1464.

65. Melby-Lervåg, M., Lyster, S.-A. H., & Hulme, C. (2012). Phonological skills and their role in learning to read: a meta-analytic review. Psychological Bulletin, 138(2), 322. doi:http://dx.doi.org/10.1037/a0026744

66. Moore, E. M., Migliorini, R., Infante, M. A., & Riley, E. P. (2014). Fetal alcohol spectrum disorders: recent neuroimaging findings. Current Developmental Disorders Reports, 1(3), 161–172.

67. Nguyen, V. T., Chong, S., Tieng, Q. M., Mardon, K., Galloway, G. J., & Kurniawan, N. D. (2017). Radiological studies of fetal alcohol spectrum disorders in humans and animal models: An updated comprehensive review. Magnetic Resonance Imaging, 43, 10–26.

68. Nichols, T. E., & Holmes, A. P. (2002). Nonparametric permutation tests for functional neuroimaging: a primer with examples. Human Brain Mapping, 15(1), 1–25.

69. O’Leary, C. M., Taylor, C., Zubrick, S. R., Kurinczuk, J. J., & Bower, C. (2013). Prenatal alcohol exposure and educational achievement in children aged 8–9 years. Pediatrics, 132(2), e468–e475.

70. Ozernov-Palchik, O., & Gaab, N. (2016). Tackling the ‘dyslexia paradox’: reading brain and behavior for early markers of developmental dyslexia. Wiley Interdisciplinary Reviews: Cognitive Science, 7(2), 156–176. doi:http://dx.doi.org/10.1002/wcs.1383

71. Pelham Jr, W. E., Gnagy, E. M., Greenslade, K. E., & Milich, R. (1992). Teacher ratings of DSM-III-R symptoms for the disruptive behavior disorders. Journal of the American Academy of Child & Adolescent Psychiatry, 31(2), 210–218.

72. Pugh, K. R., Mencl, W. E., Jenner, A. R., Katz, L., Frost, S. J., Lee, J. R., … Shaywitz, B. A. (2000). Functional neuroimaging studies of reading and reading disability (developmental dyslexia). Mental Retardation and Developmental Disabilities Research Reviews, 6(3), 207–213.

73. Pugh, K. R., Mencl, W. E., Jenner, A. R., Katz, L., Frost, S. J., Lee, J. R., … Shaywitz, B. A. (2001). Neurobiological studies of reading and reading disability. Journal of Communication Disorders, 34(6), 479–492. doi:https://doi.org/10.1016/S0021-9924(01)00060-0

74. Raschle, N. M., Stering, P. L., Meissner, S. N., & Gaab, N. (2013). Altered neuronal response during rapid auditory processing and its relation to phonological processing in prereading children at familial risk for dyslexia. Cerebral Cortex, 24(9), 2489–2501.

75. Raschle, N. M., Zuk, J., & Gaab, N. (2012). Functional characteristics of developmental dyslexia in left-hemispheric posterior brain regions predate reading onset. Proceedings of the National Academy of Sciences, 109(6), 2156–2161. doi:https://doi.org/10.1073/pnas.1107721109

76. Richlan, F., Kronbichler, M., & Wimmer, H. (2009). Functional abnormalities in the dyslexic brain: A quantitative meta-analysis of neuroimaging studies. Human Brain Mapping, 30(10), 3299–3308.

77. Richlan, F., Kronbichler, M., & Wimmer, H. (2011). Meta-analyzing brain dysfunctions in dyslexic children and adults. NeuroImage, 56(3), 1735–1742. doi:https://doi.org/10.1016/j.neuroimage.2011.02.040

78. Richlan, F., Kronbichler, M., & Wimmer, H. (2013). Structural abnormalities in the dyslexic brain: a meta-analysis of voxel-based morphometry studies. Human Brain Mapping, 34(11), 3055–3065. doi:http://dx.doi.org/10.1002/hbm.22127

79. Rimrodt, S., Clements-Stephens, A., Pugh, K., Courtney, S., Gaur, P., Pekar, J., & Cutting, L. (2009). Functional MRI of sentence comprehension in children with dyslexia: beyond word recognition. Cerebral Cortex, 19(2), 402–413.

80. Robertson, F. C., Narr, K. L., Molteno, C. D., Jacobson, J. L., Jacobson, S. W., & Meintjes, E. M. (2015). Prenatal alcohol exposure is associated with regionally thinner cortex during the preadolescent period. Cerebral Cortex, 26(7), 3083–3095.

81. Rodd, J. M., Vitello, S., Woollams, A. M., & Adank, P. (2015). Localising semantic and syntactic processing in spoken and written language comprehension: An Activation Likelihood Estimation meta-analysis. Brain and Language, 141, 89–102.

82. Roussotte, F. F., Bramen, J. E., Nunez, S. C., Quandt, L. C., Smith, L., O’Connor, M. J., … Sowell, E. R. (2011). Abnormal brain activation during working memory in children with prenatal exposure to drugs of abuse: the effects of methamphetamine, alcohol, and polydrug exposure. NeuroImage, 54(4), 3067–3075.

83. Sampson, P. D., Streissguth, A. P., Barr, H. M., & Bookstein, F. L. (1989). Neurobehavioral effects of prenatal alcohol: Part II. Partial least squares analysis. Neurotoxicology and Teratology, 11(5), 477–491.

84. Santhanam, P., Li, Z., Hu, X., Lynch, M. E., & Coles, C. D. (2009). Effects of prenatal alcohol exposure on brain activation during an arithmetic task: an fMRI study. Alcoholism: Clinical and Experimental Research, 33(11), 1901–1908.

85. Schlaggar, B. L., & McCandliss, B. D. (2007). Development of neural systems for reading. Annual Review of Neuroscience, 30, 475–503.

86. Sowell, E. R., Johnson, A., Kan, E., Lu, L. H., Van Horn, J. D., Toga, A. W., … Bookheimer, S. Y. (2008). Mapping white matter integrity and neurobehavioral correlates in children with fetal alcohol spectrum disorders. Journal of Neuroscience, 28(6), 1313–1319.

87. Sowell, E. R., Lu, L. H., O’Hare, E. D., McCourt, S. T., Mattson, S. N., O’Connor, M. J., & Bookheimer, S. Y. (2007). Functional magnetic resonance imaging of verbal learning in children with heavy prenatal alcohol exposure. Neuroreport, 18(7), 635–639.

88. Sowell, E. R., Mattson, S. N., Kan, E., Thompson, P. M., Riley, E. P., & Toga, A. W. (2008). Abnormal cortical thickness and brain–behavior correlation patterns in individuals with heavy prenatal alcohol exposure. Cerebral Cortex, 18(1), 136–144.

89. Sowell, E. R., Thompson, P. M., Mattson, S. N., Tessner, K. D., Jernigan, T. L., Riley, E. P., & Toga, A. W. (2001). Voxel-based morphometric analyses of the brain in children and adolescents prenatally exposed to alcohol. Neuroreport, 12(3), 515–523.

90. Sowell, E. R., Thompson, P. M., Mattson, S. N., Tessner, K. D., Jernigan, T. L., Riley, E. P., & Toga, A. W. (2002). Regional brain shape abnormalities persist into adolescence after heavy prenatal alcohol exposure. Cerebral Cortex, 12(8), 856–865.

91. Sowell, E. R., Thompson, P. M., Peterson, B. S., Mattson, S. N., Welcome, S. E., Henkenius, A. L., … Toga, A. W. (2002). Mapping cortical gray matter asymmetry patterns in adolescents with heavy prenatal alcohol exposure. NeuroImage, 17(4), 1807–1819.

92. Streissguth, A. P., Barr, H. M., & Sampson, P. D. (1990). Moderate prenatal alcohol exposure: effects on child IQ and learning problems at age 7 1/2 years. Alcoholism: Clinical and Experimental Research, 14(5), 662–669.

93. Streissguth, A. P., Sampson, P. D., Olson, H. C., Bookstein, F. L., Barr, H. M., Scott, M., … Mirsky, A. F. (1994). Maternal drinking during pregnancy: Attention and short-term memory in 14-year-old offspring—a longitudinal prospective study. Alcoholism: Clinical and Experimental Research, 18(1), 202–218.

94. Sullivan, E. V., Moore, E. M., Lane, B., Pohl, K. M., Riley, E. P., & Pfefferbaum, A. (2020). Graded Cerebellar Lobular Volume Deficits in Adolescents and Young Adults with Fetal Alcohol Spectrum Disorders (FASD). Cerebral Cortex, 30(9), 4729–4746.

95. Suttie, M., Foroud, T., Wetherill, L., Jacobson, J. L., Molteno, C. D., Meintjes, E. M., … Riley, E. P. (2013). Facial dysmorphism across the fetal alcohol spectrum. Pediatrics, 131(3), e779–e788.

96. Torgesen, J. K., Rashotte, C. A., & Wagner, R. K. (1999). TOWRE: Test of word reading efficiency: Pro-ed Austin, TX.

97. Treit, S., Chen, Z., Zhou, D., Baugh, L., Rasmussen, C., Andrew, G., … Beaulieu, C. (2017). Sexual dimorphism of volume reduction but not cognitive deficit in fetal alcohol spectrum disorders: A combined diffusion tensor imaging, cortical thickness and brain volume study. NeuroImage: Clinical, 15, 284–297.

98. van der Kouwe, A. J., Benner, T., Salat, D. H., & Fischl, B. (2008). Brain morphometry with multiecho MPRAGE. NeuroImage, 40(2), 559–569.

99. Vanderauwera, J., De Vos, A., Forkel, S. J., Catani, M., Wouters, J., Vandermosten, M., & Ghesquière, P. (2018). Neural organization of ventral white matter tracts parallels the initial steps of reading development: A DTI tractography study. Brain and Language, 183, 32–40.

100. Vandermosten, M., Poelmans, H., Sunaert, S., Ghesquière, P., & Wouters, J. (2013). White matter lateralization and interhemispheric coherence to auditory modulations in normal reading and dyslexic adults. Neuropsychologia, 51(11), 2087–2099.

101. Vinckier, F., Dehaene, S., Jobert, A., Dubus, J. P., Sigman, M., & Cohen, L. (2007). Hierarchical coding of letter strings in the ventral stream: dissecting the inner organization of the visual word-form system. Neuron, 55(1), 143–156.

102. Wagner, R. K., & Torgesen, J. K. (1987). The nature of phonological processing and its causal role in the acquisition of reading skills. Psychological Bulletin, 101(2), 192.

103. Wang, Y., Mauer, M. V., Raney, T., Peysakhovich, B., Becker, B. L., Sliva, D. D., & Gaab, N. (2016). Development of tract-specific white matter pathways during early reading development in at-risk children and typical controls. Cerebral Cortex, 27(4), 2469–2485.

104. Wechsler, D. (2011). WASI-II: *Wechsler abbreviated scale of intelligence*: PsychCorp.

105. Wozniak, J. R., & Muetzel, R. L. (2011). What does diffusion tensor imaging reveal about the brain and cognition in fetal alcohol spectrum disorders? Neuropsychology Review, 21(2), 133–147.

106. Wozniak, J. R., Riley, E. P., & Charness, M. E. (2019). Clinical presentation, diagnosis, and management of fetal alcohol spectrum disorder. The Lancet Neurology, 18(8), 760–770.

107. Yang, Y., Phillips, O. R., Kan, E., Sulik, K. K., Mattson, S. N., Riley, E. P., … O’Connor, M. J. (2012). Callosal thickness reductions relate to facial dysmorphology in fetal alcohol spectrum disorders. Alcoholism: Clinical and Experimental Research, 36(5), 798–806.

108. Yang, Y., Roussotte, F., Kan, E., Sulik, K. K., Mattson, S. N., Riley, E. P., … O’Connor, M. J. (2012). Abnormal cortical thickness alterations in fetal alcohol spectrum disorders and their relationships with facial dysmorphology. Cerebral Cortex, 22(5), 1170–1179.

109. Yeatman, J. D., Dougherty, R. F., Ben-Shachar, M., & Wandell, B. A. (2012). Development of white matter and reading skills. Proceedings of the National Academy of Sciences, 109(44), E3045–E3053.

110. Yeatman, J. D., Dougherty, R. F., Myall, N. J., Wandell, B. A., & Feldman, H. M. (2012). Tract profiles of white matter properties: automating fiber-tract quantification. PloS One, 7(11), e49790.

111. Yu, X., Raney, T., Perdue, M. V., Zuk, J., Ozernov-Palchik, O., Becker, B. L., … Gaab, N. (2018). Emergence of the neural network underlying phonological processing from the prereading to the emergent reading stage: A longitudinal study. Human Brain Mapping, 39(5), 2047–2063.

112. Yu, X., Zuk, J., Perdue, M. V., Ozernov-Palchik, O., Raney, T., Beach, S. D., … Gaab, N. (2020). Putative protective neural mechanisms in prereaders with a family history of dyslexia who subsequently develop typical reading skills. Human Brain Mapping, 41(10), 2827–2845.

